# Integrated Stress Response signaling acts as a metabolic sensor in fat tissues to regulate oocyte maturation and ovulation

**DOI:** 10.1101/2023.02.27.530289

**Authors:** Lydia Grmai, Manuel Michaca, Emily Lackner, Narayanan Nampoothiri V.P., Deepika Vasudevan

## Abstract

Reproduction is an energy-intensive process requiring systemic coordination. However, the inter-organ signaling mechanisms that relay nutrient status to modulate reproductive output are poorly understood. Here, we use *Drosophila melanogaster* as a model to establish the Integrated Stress response (ISR) transcription factor, Atf4, as a fat tissue metabolic sensor which instructs oogenesis. We demonstrate that Atf4 regulates the lipase Brummer to mediate yolk lipoprotein synthesis in the fat body. Depletion of *Atf4* in the fat body also blunts oogenesis recovery after amino acid deprivation and re-feeding, suggestive of a nutrient sensing role for Atf4. We also discovered that Atf4 promotes secretion of a fat body-derived neuropeptide, CNMamide, which modulates neural circuits that promote egg-laying behavior (ovulation). Thus, we posit that ISR signaling in fat tissue acts as a “metabolic sensor” that instructs female reproduction: directly, by impacting yolk lipoprotein production and follicle maturation, and systemically, by regulating ovulation.

## Introduction

The fundamental behaviors of an organism, such as feeding and reproduction, are informed by its metabolic status. Reproduction relies on nutrient availability to support the high energetic cost of gametogenesis and associated reproductive behaviors. Across multicellular organisms, the fat tissue serves as a lipid storage organ and acts as a signaling hub to peripheral organs to communicate nutrient status^1^. Consequently, fat tissue homeostasis broadly influences reproductive capacity; defects in such homeostasis due to insufficient or excess dietary lipids result in decreased fertility^2,3^. However, the inter-organ signaling mechanisms which relay nutrient status from fat tissues to inform the rate of reproductive output in the ovary are still not fully understood.

Highly metabolic tissues, such as fat and liver, have been reported to rely on stress response pathways to maintain homeostasis^4–6^. Specifically, constitutive activity of the Integrated Stress Response (ISR), an evolutionarily conserved pathway that relies on stress-sensing kinases, has been observed in adipocytes and hepatocytes^5,7^. There are four known ISR kinases-GCN2, PERK, HRI and PKR-that all signal through stress-response transcription factors when activated by endogenous or external stressors^8^. The best studied transcription factor downstream of the ISR kinases is Activating transcription factor 4 (ATF4)^8^. The importance of ISR signaling in metabolic tissues is highlighted by patient mutations in ISR effectors and loss of function studies in model organisms. PERK mutations in humans cause Wolcott-Rallison syndrome characterized by neonatal diabetes^9,10^, and loss of PERK signaling in mouse models results in dysregulated liver glycogen content and increased hepatocyte death in high sugar dietary conditions^11,12^. In mice and fruit flies (*Drosophila melanogaster*), *Atf4* mutant animals show lower body fat content, and *ATF4*^*-/-*^ mice show greater resistance to fatty liver under high dietary intake conditions^4^.

We and others have shown PERK and GCN2 to be constitutively active in fat tissues in mice and in *Drosophila*^7,13,14^. Similar to *Atf4*, loss of *Perk* or *Gcn2* results in dysregulation of fatty acid homeostasis, which is further exacerbated by dietary restriction or excess lipid or sugar intake^11,13^. While the primary effects of the loss of ISR factors on fat tissues have been well studied, how ISR-mediated fat tissue homeostasis impacts peripheral tissue function has not been reported. Here, we use the *Drosophila* model to investigate how ISR signaling in fat tissue (called the ‘fat body’) affects female reproduction. The fat body is a highly metabolic tissue comprised by multiple cell types that together execute vertebrate liver and adipose tissue functions, including fat storage, detoxification and immune response^1^. A prominent role for the female fat body is the synthesis and trafficking of yolk lipoprotein to oocytes, which is critical for oocyte maturation^15^.

The *Drosophila* ovary is organized as long chains of developing follicles called ovarioles^16^ **(Fig. 1A)**. Germline and somatic stem cells reside at the anterior apex and undergo differentiation along the ovariole in individual follicles. The germline stem cells differentiate to give rise to sixteen-cell germ cell ‘cysts’, with one designated to be the oocyte and fifteen supporting nurse cells. Concomitantly differentiating somatic cells envelop the germ cyst to form a follicle. Each follicle undergoes 14 stages of oogenesis, culminating in a mature oocyte **(Fig. 1A)**. From stages 8-14, the oocyte accumulates yolk lipoprotein, which is synthesized both by follicle cells within the ovary and fat body surrounding the ovary^17^. We previously reported that loss of *Atf4*, encoded by *cryptocephal (crc)* in *Drosophila*, resulted in mid-oogenesis arrest accompanied by follicle death^18^. In this study, we performed tissue-specific depletion of *Atf4* to determine the basis of these oogenesis defects.

**Figure 1.**
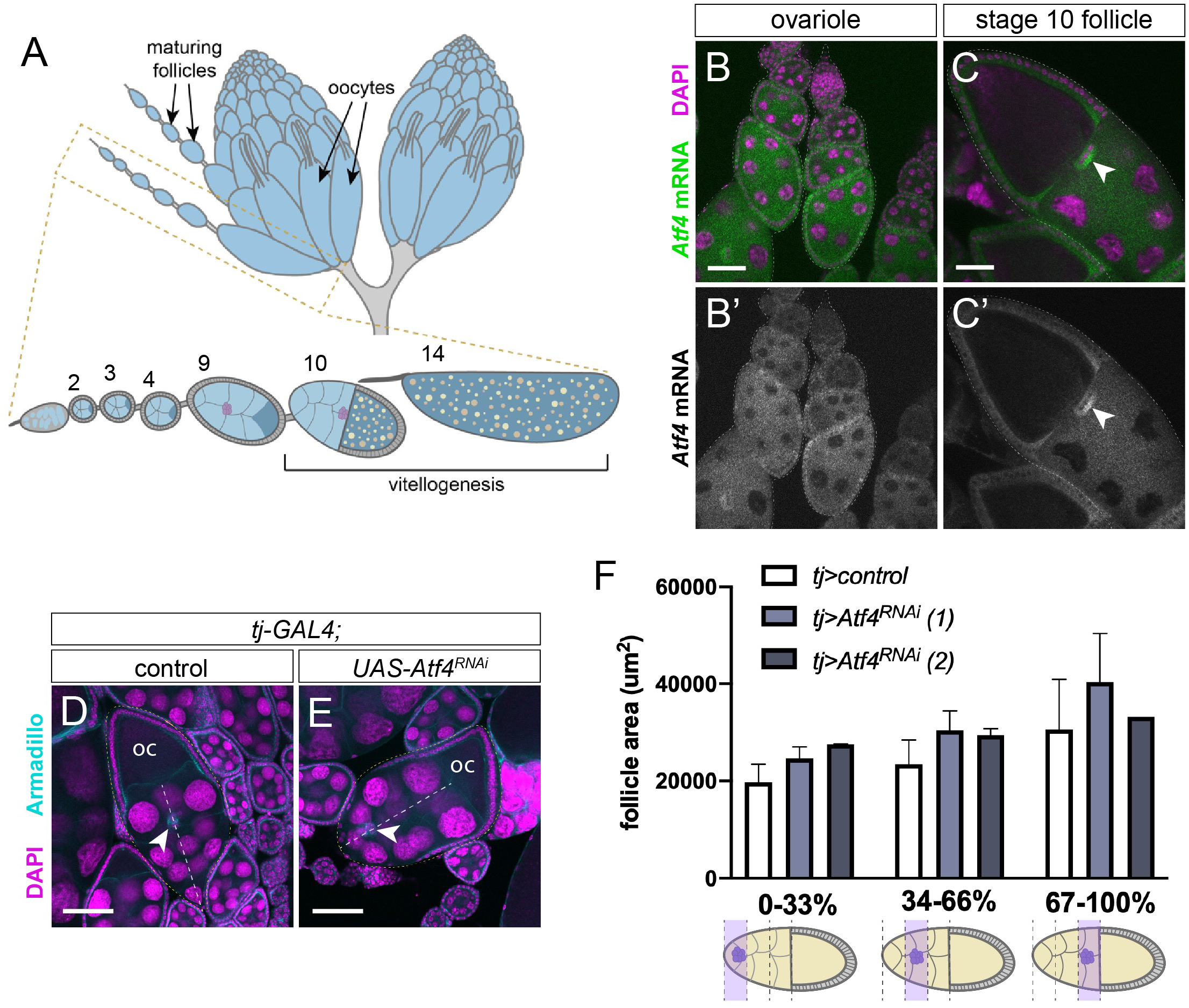
*Atf4* is required in the ovary for oogenesis. (A) Schematic of adult *Drosophila* ovaries showing clusters of ovarioles containing maturing follicles. Diagram on the bottom shows an individual ovariole with various follicle stages. Vitellogenesis begins during mid-oogenesis, when yolk lipoproteins (orange spots) are trafficked from somatic cells (light blue) into maturing oocytes (dark blue). Border cells are shown stage 9 (purple cells) and complete their migration at the anterior end of the stage 10 follicle. (B-C) Visualization of *Atf4* mRNA (green) in adult ovarioles (B, B’) and stage 10 follicles (C, C’) by FISH. Scale bars: 50 µm. (D-E) Representative confocal images of border cells in stage 10 follicles from control (D) and *tj>Atf4*^*RNAi*^ (E) ovaries isolated from 5-day old adult females. Follicles of roughly equal size from each genotype are indicated in yellow dotted outline; white dotted line indicates border cell migration path, beginning at the anterior end of the follicle and terminating at the oocyte (oc). Border cells are identified with a white arrow. In control follicle (D), border cell is approximately 75% migrated, while border cells lacking *Atf4* (E) have only migrated approximately 24% at this follicle size. See **Fig. S1E** for calculation methods. Scale bars: 100 µm. (F) Graph plotting percent border cell migration versus area of the containing follicle in *control* (blue) or *tj>Atf4*^*RNAi*^ (gray/orange lines represent two independent *Atf4*^*RNAi*^ lines). In microscopy figures, DAPI marks nuclei in magenta. Schematic underneath graph shows extent of border cell migration quantified. See **Fig. S1E** for calculation methods,

## Results

### ISR factors are required in the ovary for oogenesis

We previously showed that global *Atf4*-mutant animals exhibit oogenesis defects, including mid-oogenesis arrest and increased follicle death^18^. To determine if these oogenesis defects were ovary-autonomous, we had examined Atf4 expression in the ovary. We observed that an Atf4-GFP fusion protein encoded by a protein trap allele (*Atf4*^*GFSTF*^) was not detected in adult ovaries^18^. To test whether an undetectably low level of Atf4 expression might play a tissue-autonomous role in the ovary, we probed expression of *Atf4* mRNA using fluorescent *in situ* hybridization (FISH). We found *Atf4* mRNA in differentiating germline cells and somatic follicle cells beginning around stage 5 through stage 10 **(Fig. 1B-C)**.

Since *Atf4* mRNA was present in multiple ovary cell types, we sought to test whether depletion of *Atf4* in the ovary results in follicle death or mid-oogenesis arrest similar to that seen in global *Atf4* mutants^18^. To do so, we utilized the GAL4/UAS binary expression system^19^ to deplete *Atf4* specifically in either germline or somatic cells. We detected follicle death either by the presence of cleaved caspase staining (Dcp1 in *Drosophila*) or by observing for germ cell nuclear fragmentation, which is indicative of follicle death. We found that RNAi depletion of *Atf4* using the germline driver *nanos-GAL4* did not result in increased follicle death in comparison to ovaries from control animals (**Fig. S1A**). In addition, we found no evidence of increased oogenesis arrest, as seen by the number of follicles in mid-oogenesis per ovary (**Fig. S1B**). We next used the somatic gonad driver *tj-GAL4* to deplete *Atf4* and found that ovaries lacking *Atf4* in the somatic gonad showed no significant increase in follicle death or oogenesis arrest when compared to control ovaries **(Fig. S1C-D)**. Thus, our data indicate that the oogenesis defects observed in *Atf4* mutant animals^18^ are not due to an autonomous requirement for *Atf4* in the ovary.

Notably, our FISH analysis also revealed a striking enrichment of *Atf4* mRNA in border cells (**white arrow in Fig. 1B-C**). The border cells are a cluster of 8-10 follicular epithelial cells which delaminate from the anterior end of the follicle and migrate to the anterior end of the oocyte (**Fig. 1A**)^20^. Collectively, the border cells establish oocyte polarity and form egg structures that are critical for embryonic survival, such as the dorsal appendage^21^. While loss of *Atf4* in the somatic gonad did not result in increased follicle death, we observed a delay in border cell migration in these follicles. We quantified the extent of border cell migration by measuring follicle size and percent border cell migration across stage 9-10 follicles (**Fig. S1E**)^22^. The average area of follicles with border cells in the first, second, or final third of migration was then plotted for each genotype, similar to previously published characterizations of border cell migration^23^. Using such analysis, and the border cell marker Armadillo, we noticed a delay in border cell migration in *tj>Atf4*^*RNAi*^ follicles compared with controls (**Fig. 1D-F**). We validated these results using a second independent RNAi line targeting *Atf4*, which showed a similar delay in border cell migration (**Fig. 1F**). Similar results were simultaneously obtained by Dr. Jocelyn McDonald’s group, who has characterized this border cell migration defect extensively in an accompanying manuscript.

We next turned to testing whether ISR signaling outside of the ovary is responsible for the follicle death and mid-oogenesis arrest phenotypes observed in *Atf4* mutants. Given the notable role of ISR signaling in fat tissue homeostasis^5^, we performed RNAi-mediated knockdown of *Atf4* using the fat body-specific driver *3.1Lsp2-GAL4*^24^. Fat body depletion of *Atf4* caused follicle death and mid-oogenesis arrest reminiscent of that observed in *Atf4* mutant ovaries^18^ (**Fig. 2A-D**). Taken together, these results led us to pursue a non-autonomous role for fat body Atf4 signaling in oogenesis, which we describe below.

**Figure 2.**
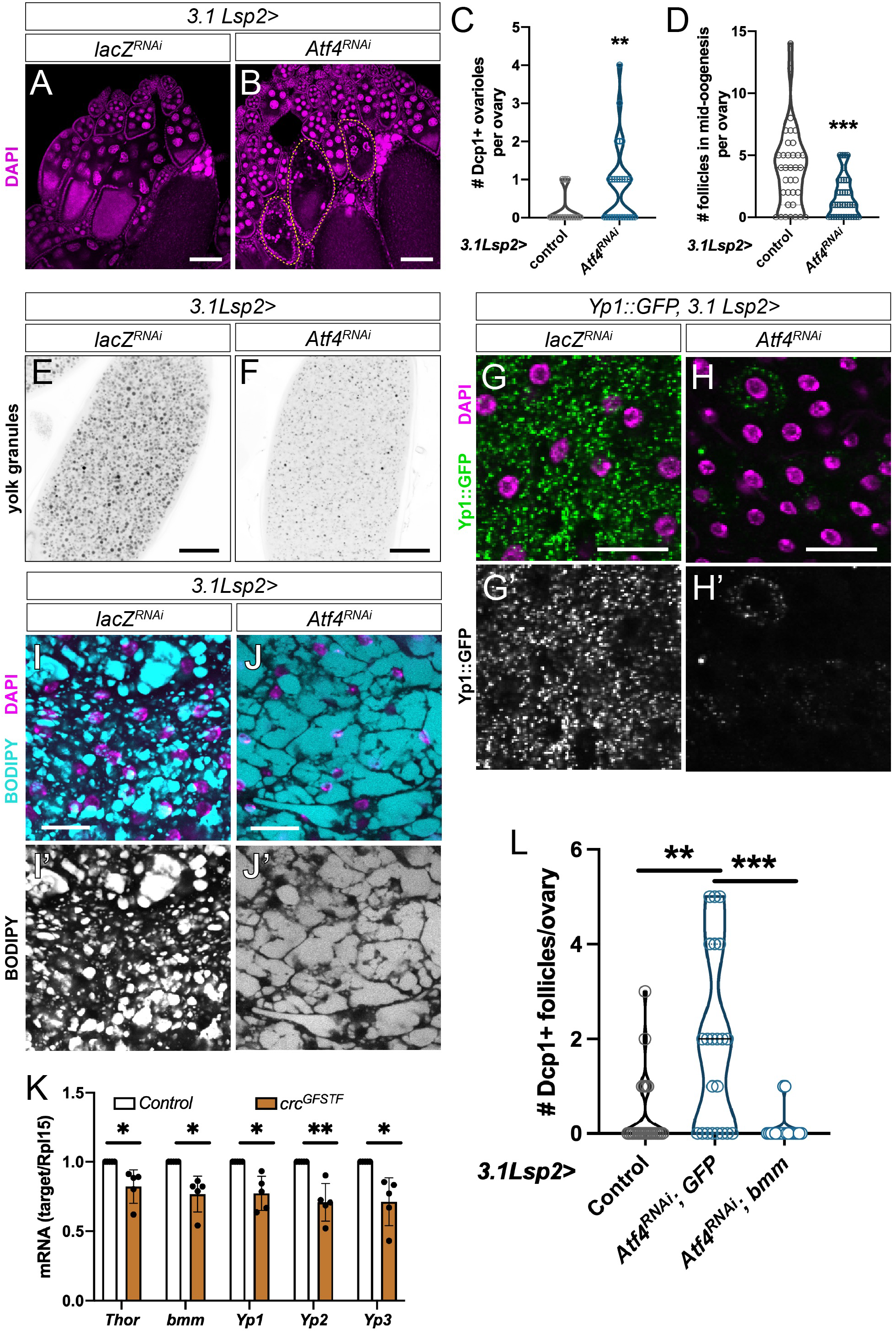
ISR signaling promotes proper yolk protein secretion from the fat body. (A-B) Representative images of adult ovaries following control (*lacZ*, A) or *Atf4* (B) depletion from the fat body using *3.1Lsp2-GAL4*. Ovarioles marked with a dotted outline show germ cell nuclear fragmentation indicative of dying follicles. (C) Quantification of follicle death from A-B as seen by cleaved caspase (Dcp1) staining. Death was quantified on the y-axis as the number of ovarioles containing a dying follicle per ovary in each genotype. (D) Quantification of oogenesis arrest from A-B, reported as the number of follicles in mid-oogenesis (stages 6-10) per ovary. (E-F) Representative confocal images of stage 14 oocytes from *3.1Lsp2>lacZ*^*RNAi*^ (E) and *>Atf4*^*RNAi*^ (F) females. Yolk granules were visualized by auto-fluorescence with 405nm laser, yolk granules appear as black/gray circles. Scale bar: 200 µm. (G-H) Representative confocal images of Yp1::GFP (green) expression in adult fat body from *3.1Lsp2>lacZ*^*RNAi*^ (G) and *Atf4*^*RNAi*^ (H) females. Scale bar: 50 µm. (I-J) Representative confocal images of neutral lipid staining (BODIPY, cyan) in adult fat body from *3.1Lsp2>lacZ*^*RNAi*^ (I) and *Atf4*^*RNAi*^ (J) females. Scale bar: 50 µm. (K) qPCR analysis of transcript abundance in fat bodies from control (*yw*) versus *Atf4* hypomorphic mutants (*crc*^*GFSTF*^) for indicated transcripts normalized to *Rpl15* as a house-keeping gene. Values are reported as the average of five biological replicates. (L) Quantification of follicle death (as determined by Dcp1 staining) in control ovaries compared with loss of *Atf4* (*3.1Lsp2>Atf4*^*RNAi*^; *>GFP*) and rescue by *bmm* co-expression (*3.1Lsp2>Atf4*^*RNAi*^, *>bmm*). In confocal images, DAPI (magenta) labels DNA. Here and in future figures, statistical analyses were performed using a two-tailed unpaired Student’s t-test with Welch’s correction. Asterisks indicate statistical significance as follows: *p<0.05; **p<0.01; ***p<0.001; ****p<0.0001; ns = not significant.

### Atf4 promotes yolk lipoprotein secretion from the fat body

We first sought to determine possible causes of the follicle death observed upon loss of Atf4 signaling in the fat body. Follicle death in *3.1Lsp2>Atf4*^*RNAi*^ was frequently observed in mid-oogenesis (**Fig. 2B-C**). A prominent checkpoint in mid-oogenesis is yolk and lipid uptake (vitellogenesis), which occurs in stage 8-14 follicles^17^ (**Fig. 1A)**. In *Drosophila*, yolk proteins are synthesized in both the somatic gonad and the fat body^15^. We found that fat body-specific depletion of *Atf4* resulted in substantially less yolk granule accumulation in stage 14 oocytes compared to controls **(Fig. 2E-F)**. Thus, we hypothesized that loss of ISR factors in the fat body compromises yolk protein trafficking to the ovary, which results in follicle death by disrupting vitellogenesis.

Yolk proteins are trafficked from the fat body to the ovary as lipoprotein vesicles, which contain a lipid core surrounded by the lipoproteins Yp1, Yp2, and Yp3^25^. The lipid core of lipoproteins is derived from lipid droplets, which are extensions of the endoplasmic reticulum (ER) that bud off to form protein-coated vesicles^26^. We tested whether there was an overall decrease in yolk lipoprotein production by visualizing expression of a Yolk protein 1 (Yp1)-GFP fusion protein (Yp1::GFP)^27^ in the fat body. Knockdown of *Atf4* in the fat body showed a dramatic reduction of Yp1::GFP in adipocytes **(Fig. 2G-H)**, suggesting that loss of *Atf4* resulted in impaired yolk lipoprotein synthesis in the fat body. Given the prominent role for Atf4 in lipid homeostasis^4,28^, we assayed for changes in lipid droplet formation in the fat body with loss of *Atf4*. Staining with the neutral lipid dye BODIPY showed that control fat body contained an abundance of lipid droplets that characteristically organized into vesicles of varying sizes **(Fig. 2I)**. In contrast, the BODIPY staining in *Atf4*-depleted fat body appeared diffuse, with a marked absence of discrete vesicular structures **(Fig. 2J)**.

We next sought to determine the underlying mechanism by which Atf4 affects lipoprotein synthesis by probing for *Atf4* occupancy on a broad list of lipid homeostasis genes^29^ using the ENCODE ChIP-Seq data^30,31^. This dataset showed enrichment of Atf4 binding at the *Thor* locus, a known transcriptional target of Atf4^7^ (**Fig. S2A, 2K**), suggesting that these data can predict direct Atf4 targets. Our analysis showed several Atf4 binding sites within the gene locus of *brummer* (*bmm*) **(Fig. S2A)**, which encodes a triglyceride lipase orthologous to mammalian adipose triglyceride lipase (ATGL)^32^. Brummer localizes to the surface of lipid droplets to mobilize triacylglycerol therein^32^, which is a key step in the formation of the lipid core in lipoproteins. These observations prompted us to examine whether *bmm* is transcriptionally regulated by *Atf4* in the fat body. Since the adult fat body is proximal to multiple cell types that do not express *3.1Lsp2-GAL4*, which could confound our analysis^1^, we instead performed qPCR analysis on global *Atf4* mutants using a hypomorphic allele (*crc*^*GFSTF*^). Our analysis showed a marked decrease of *bmm* transcript in *Atf4* mutants in comparison to control animals (**Fig. 2K**), suggesting that *bmm* may indeed be a direct transcriptional target of Atf4. RNAi-mediated depletion of *bmm* from the fat body increased the size of lipid droplets, though to a lesser extent than with fat-specific *Atf4* depletion (**Fig. S2B-C**). We next tested whether restoring *bmm* expression was sufficient to rescue follicle death seen with loss of *Atf4* in the fat body (**Fig. 2A-C**). Indeed, we found that co-expressing *bmm* in the fat body reduced the number of dying follicles in *3.1Lsp2>Atf4*^*RNAi*^ animals (**Fig. 2L**).

We reasoned that yolk lipoprotein defects in *3.1Lsp2>Atf4*^*RNAi*^ animals could also be due to reduced yolk protein (*Yp1-3*) transcription in the fat body. Our data showed a substantial decrease in the transcript levels of *Yp1, Yp2*, and *Yp3* (**Fig. 2K**). However, analysis of the ENCODE project^30,31^ ChIP-seq data revealed no Atf4 occupancy at the *Yp1, Yp2*, or *Yp3* loci (**Fig. S2A**), which suggests that these genes are likely not direct transcriptional targets of Atf4.

Taken together, these data demonstrate a role for Atf4 in yolk lipoprotein synthesis and assembly, underscoring the importance of fat body-derived yolk lipoproteins in follicle maturation. Further, these data support our hypothesis that fat body ISR signaling supports oogenesis in part by ensuring sufficient yolk lipoprotein production for deposition into maturing oocytes.

### Atf4 in the fat body acts as a nutrient sensor to inform oogenesis

Nutrient deprivation, which is known to induce oogenesis arrest and promote follicle death in a reversible manner^33,34^, induces Atf4 signaling^8^. Thus, we asked whether oogenesis arrest during nutrient deprivation is mediated by Atf4. To do this, we performed an amino acid deprivation assay (**Fig. 3A, top diagram**), wherein mated females were raised for four days on amino acid-deficient medium (starved) or nutrient-rich medium (fed). We measured the rate of oogenesis by quantifying the number of follicles per ovary that were in mid-oogenesis (stages 6-10). As expected, at the end of the starvation period, ovaries from “starved” females contained significantly fewer follicles in mid-oogenesis compared with ovaries from “fed” females in control animals (**Fig. 3B**). In addition, we saw a significant increase in follicle death in “starved” versus “fed” ovaries, as determined by Dcp1 staining (**Fig. 3C**). Such oogenesis arrest and follicle death was also observed in ovaries from starved versus fed *3.1Lsp2>Atf4*^*RNAi*^ females (**Fig. 3B**).

**Figure 3.**
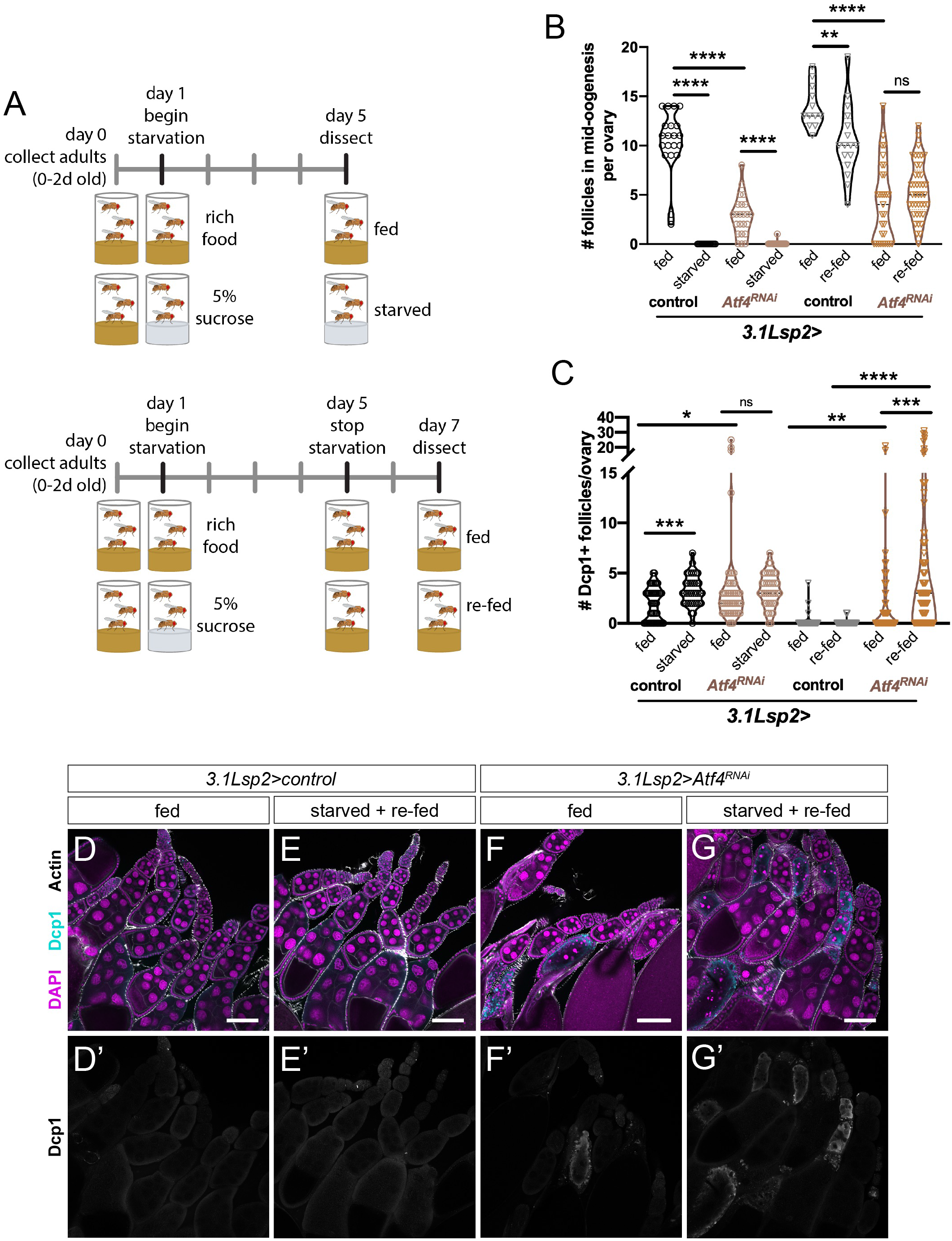
Atf4 is required for oogenesis recovery after amino acid deprivation. (A) Diagram illustrating starvation and re-feeding assay protocol. See methods and results for more details. (B) Quantification of follicles in mid-oogenesis per ovary from control and *3.1Lsp2>Atf4*^*RNAi*^ females under fed, starved, or re-fed conditions. (C) Quantification of follicle death (as determined by Dcp1 staining) per ovary from control and *3.1Lsp2>Atf4*^*RNAi*^ females under fed, starved, or re-fed conditions. (D-G) Representative confocal images of ovaries from control (D-E) and *3.1Lsp2>Atf4*^*RNAi*^ (F-G) females under fed (D,F) or starved and re-fed (E,G) conditions. DNA is labeled with DAPI (magenta). Scale bar: 100 µm. Statistical analyses were performed using a two-tailed unpaired Student’s t-test with Welch’s correction.

The oogenesis arrest induced by nutrient deprivation can typically be restored to near-normal levels by re-introduction of amino acids to the diet^33^. Consistently, we found that re-feeding control females with nutrient-rich food following the four-day starvation (**Fig. 3A, bottom diagram**) increases the number of mid-oogenesis follicles per ovary (**Fig. 3B**). Additionally, we also found that the follicle death following starvation is substantially reduced after re-feeding in control animals (**Fig. 3C-E**). Analysis of ovaries from *3.1Lsp2>Atf4*^*RNAi*^ females showed resumption of oogenesis following starvation as seen by the number of follicles in mid-oogenesis (**Fig. 3B**). However, we also saw a massive increase in the rate of follicle death in these ovaries after the re-feeding period (**Fig. 3C, F, G**), indicating impaired recovery of oogenesis after nutrient deprivation in *3.1Lsp2>Atf4*^*RNAi*^ females. From these data, we conclude that Atf4 in the fat body is required for sensing changes in nutrient availability and regulates physiological output by communicating nutrient status to the ovary.

### Fat body ISR signaling independently modulates ovulation

There are two known ISR kinases upstream of Atf4 in *Drosophila*: the ER-stress sensor Perk and amino acid sensor Gcn2. To assess which of these kinases are required for Atf4 signaling in the fat body, we used previously validated RNAi lines to deplete *Perk* or *Gcn2*^7^ in the fat body using *3.1Lsp2-GAL4*. Similar to *Atf4* knockdown, we found that knockdown of either *Perk* or *Gcn2* in the fat body resulted in increased follicle death (**Fig. 4A-D**) and mid-oogenesis arrest (**Fig. 4E**). Examination of maturing eggs from these animals also revealed a decrease in yolk granules as seen with loss of *Atf4* in the fat body (**Fig. S3A-C)**. These data suggest that both Perk and Gcn2 act upstream of Atf4 in the fat body.

**Figure 4.**
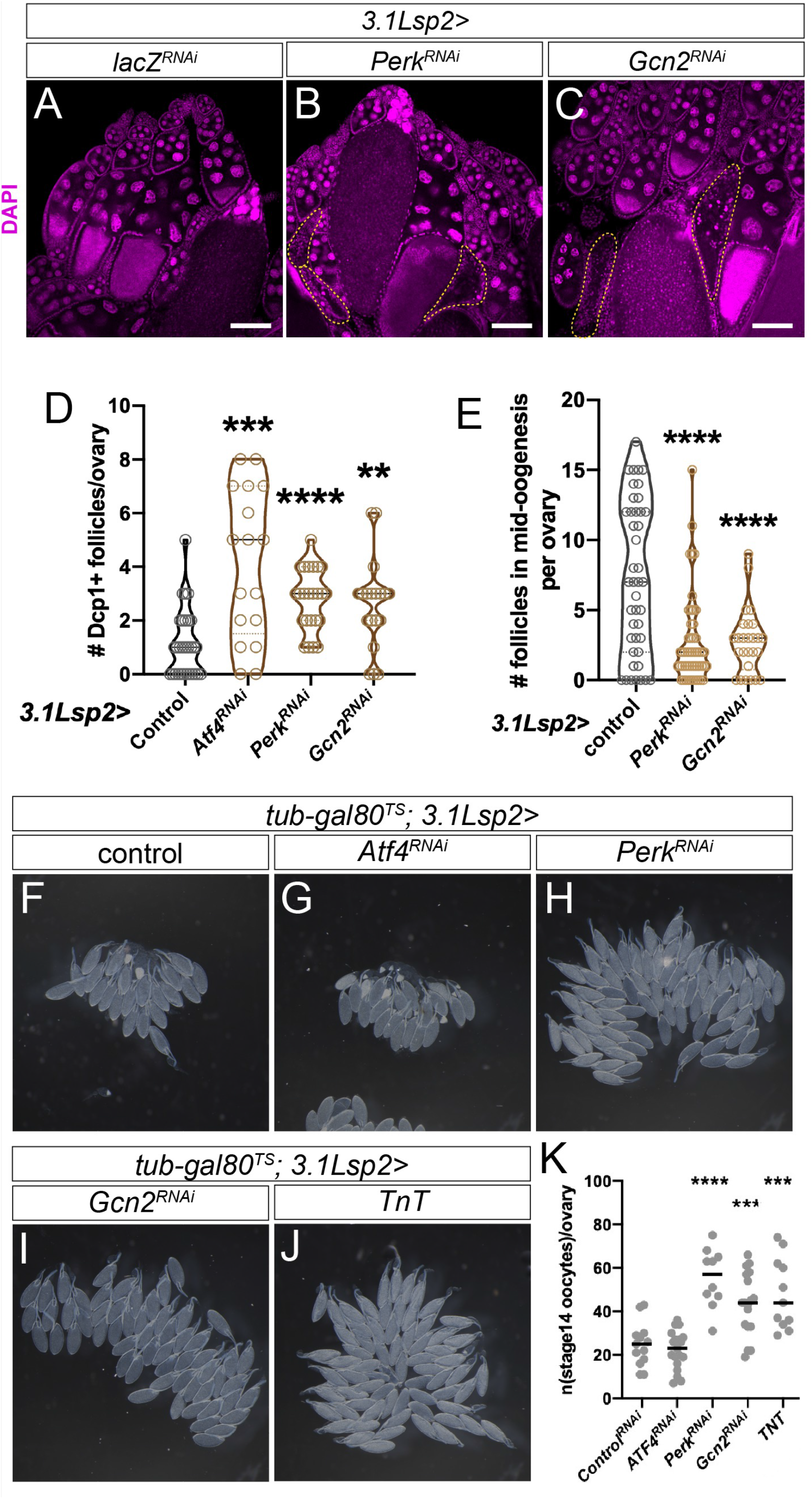
ISR signaling in the fat body modulates egg laying behavior. (A-C) Representative ovaries from *3.1Lsp2>lacZ*^*RNAi*^ (A), >*Perk*^*RNAi*^ (B), and >*Gcn2*^*RNAi*^ (C) females. Dying follicles are encircled with a yellow dotted line, identified by nuclear breakdown. DAPI is labeled in magenta. (D) Quantification of follicle death depicted in A-C as determined by Dcp1 staining. (E) Quantification of oogenesis arrest from A-C, reported as the number of follicles in mid-oogenesis (stages 6-10) per ovary. (F-J) Representative brightfield images of ovaries from *3.1Lsp2>lacZ*^*RNAi*^ (F), *Atf4*^*RNAi*^ (G), *Perk*^*RNAi*^ (H), *Gcn2*^*RNAi*^ (I), and *TnT* (J) females. Opaque oocytes located at the base of each ovary represent eggs retained in each ovary at time of dissection. (K) Quantification of egg retention phenotype shown in F-J, reported as the number of stage 14 oocytes per ovary in the indicated genotypes.

Intriguingly, in addition to increased follicle death and mid-oogenesis arrest, we also observed a paradoxical accumulation of excess oocytes in *Perk* and *Gcn2* knockdown animals (**Fig. 4F, H, I, K**). Ovaries from mated females upon fat body-specific depletion of *Perk* or *Gcn2* (*3.1Lsp2>Perk*^*RNAi*^ or *>Gcn2*^*RNAi*^) contained nearly twice as many mature oocytes per ovary (*Perk*^*RNAi*^: 55.4 ± 4.27, p<0.0001; *Gcn2*^*RNAi*^: 43.56 ± 3.39, p<0.001) than control ovaries (24.77 ± 2.78) (**Fig. 4F, H, I, K**). Ovulation is exhibited in *Drosophila* by egg-laying behavior, and defects in this process result in retention of mature eggs in the ovary^35^. While we did not observe this “egg retention” phenotype with *3.1Lsp2>Atf4*^*RNAi*^ (**Fig. 4G**; 22.26 ± 2.00), we had previously reported a ‘swollen ovary’ appearance in *Atf4* hypomorphic mutants^18^. We reasoned that since both Perk and Gcn2 signal to induce Atf4, loss of either kinase may result in weaker effects than loss of *Atf4*. This interpretation is supported by our observation that loss of *Perk* or *Gcn2* resulted in fewer Dcp1-positive follicles in comparison to loss of *Atf4* (**Fig. 4D**). We further tested this by simultaneous depletion of *Perk* and *Gcn2* in the fat body. Indeed, *3.1Lsp2>Perk*^*RNAi*^*+Gcn2*^*RNAi*^ showed no egg retention phenotype (**Fig. S3D**), similarly to *3.1Lsp2>Atf4*^*RNAi*^ ovaries. Based on this, we conclude that substantial depletion of ISR signaling (by depleting the common downstream target, Atf4) leads to severe oogenesis arrest. Such arrest masks potential ovulation defects in these animals due to an overall decrease in maturing oocytes. Consequently, a role for ISR signaling in ovulation is only revealed with partial loss of function by depleting either upstream ISR kinase, or by use of a hypomorphic *Atf4* allele.

Since we observed that ISR signaling regulates yolk lipoprotein secretion from the fat body to support oogenesis, we sought to examine whether other factors are similarly secreted from the fat body to regulate ovulation. To test this, we expressed Tetanus toxin (TnT) in the fat body to block secretion mediated by vesicular exocytosis^36^. Expression of *TnT* using *3.1Lsp2-GAL4* recapitulated the “egg retention” phenotype we observed with *3.1Lsp2>Perk*^*RNAi*^ or *>Gcn2*^*RNAi*^ compared with control animals (48.09 ± 4.94) (**Fig. 4J-K**; 48.09 ± 4.94). Thus, in addition to regulating yolk lipoprotein synthesis, our genetic analysis revealed a new role for ISR signaling in ovulation, which we further explore below.

### ISR signaling in the fat body promotes ovulation via CNMa-dependent activation of sexually dimorphic neurons

Our results thus far suggest that ISR signaling in the fat body promotes ovulation via a secreted factor. In *Drosophila*, ovulation is stimulated in mated females via activation of sexually dimorphic neural circuits^37–41^. Thus, we hypothesized that the secreted factor downstream of ISR signaling is a neuropeptide, which signals to subsets of these neurons to regulate ovulation. To identify putative Atf4-regulated neuropeptides, we compiled a comprehensive list of the annotated neuropeptides in the *D. melanogaster* genome based on FlyBase categorizations (**Table S2)**. We next examined Atf4 occupancy in the genomic loci of these neuropeptides using the ENCODE ChIP-seq dataset described above^30,31^. Analysis of the overlap between these two datasets revealed four neuropeptides with evidence of Atf4 occupancy (**Table S2**, highlighted in green). We systematically depleted each of these neuropeptides in the fat body (using *3.1Lsp2-GAL4*), and quantified egg retention (**Fig. 5A, B**). We found that depletion of all four individual neuropeptides predicted to be Atf4 targets led to increased egg retention: *short neuropeptide F (sNPF), Ecdysis triggering hormone (Eth), Tachykinin (Tk)*, and *CNMamide (CNMa)* (**Fig. 5A, B**). Further qPCR analysis using *Atf4* mutants showed a decrease in the mRNA abundance of two of these: *CNMa* and *Eth* (**Fig. 5C**). *Tk* transcripts were undetectable by qPCR in both control and *Atf4* mutants, and publicly available data via FlyAtlas and modENCODE sequencing projects report no detectable *Tk* expression in larval or adult fat body. Of these, we found that fat body-specific depletion of CNMa had the largest effect on egg retention, to similar extents as seen with *3.1Lsp2>Perk*^*RNAi*^ or *>Gcn2*^*RNAi*^ (**Fig. 5B, E**; compare with **Fig. 4H, I, K**), which prompted us to further characterize the role of this neuropeptide in ovulation.

**Figure 5.**
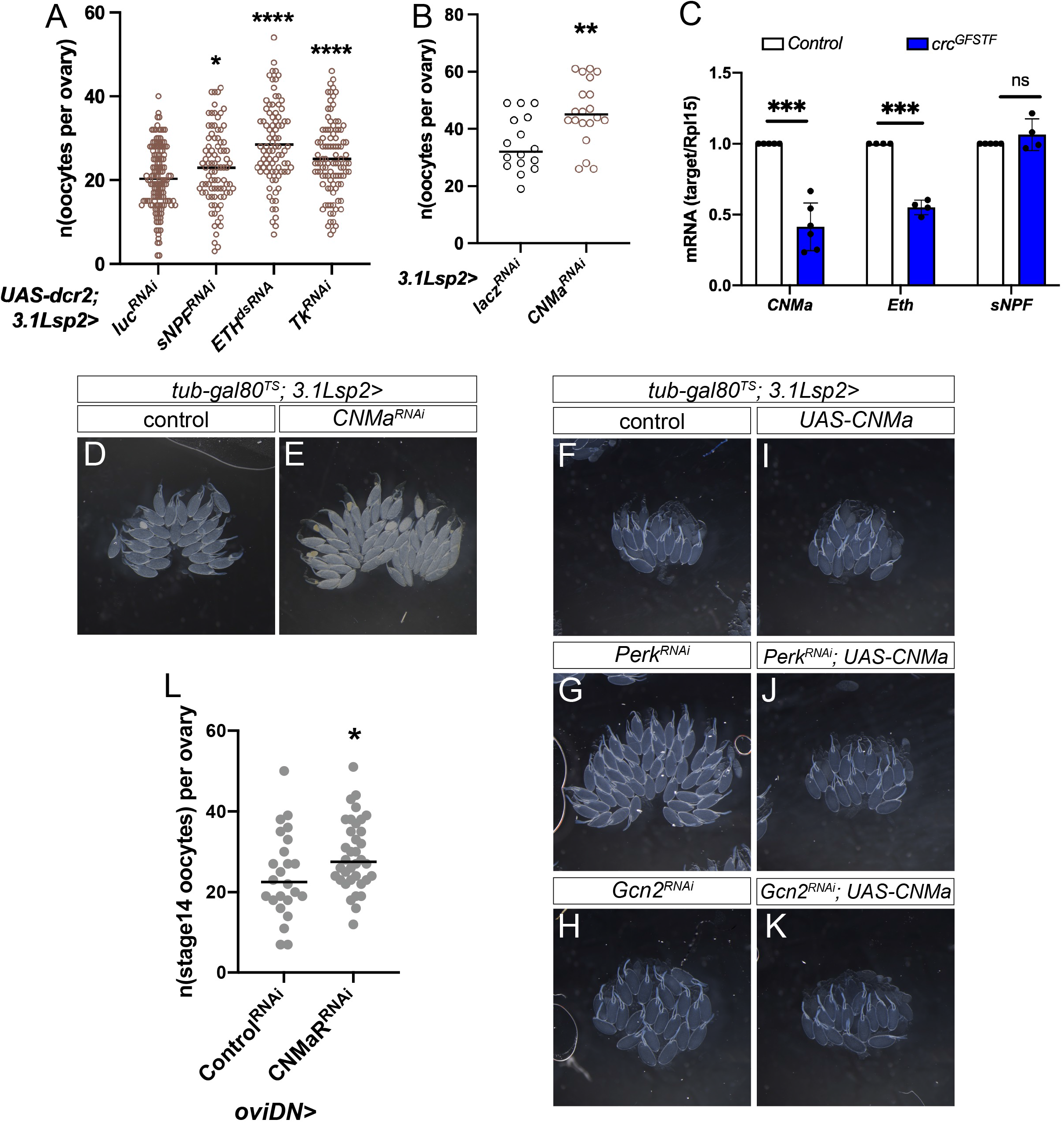
Oviposition-descending neurons (oviDNs) respond to ISR-regulated CNMa to promote ovulation. (A-B) Quantification of oocytes contained per ovary in a limited neuropeptide screen (see **Table S2** for list), wherein putative Atf4-regulated neuropeptides were individually depleted by RNAi in the fat body. Please note that RNAi lines were obtained from multiple sources, necessitating different negative controls. (C) qPCR analysis of neuropeptide transcript abundance in fat bodies from control (*yw*) versus *Atf4* hypomorphic mutants (*crc*^*GFSTF*^) for *CNMa, Eth, and sNPF* normalized to *Rpl15* as a house-keeping gene. Values are reported as the average of at least four biological replicates. Please note that *Tk* mRNA could not be detected. (D-E) Representative brightfield images of ovaries from *3.1Lsp2>control* (D) and *>CNMa*^*RNAi*^ (E) females. (F-K) Representative brightfield images of ovulation rescue by simultaneous overexpression of CNMa in the fat body. (L) Quantification of egg retention phenotype in animals where the CNMa receptor (CNMaR) is depleted in sexually dimorphic neurons using a split *oviDN-GAL4* reported as the number of stage 14 oocytes per ovary in the indicated genotypes. Error bars represent SEM.

CNMa has been previously described to be induced in enterocytes of the gut in response to amino acid deprivation in an Atf4-dependent manner^42^. Using a CNMa-GAL4 transgene^42^, we found that the *CNMa* promoter is active in both larval and adult female fat body (**Fig. S4A, B**). Consistent with our prediction that CNMa is an Atf4 target, qPCR analysis showed lower levels of *CNMa* transcripts in *Atf4* mutants (**Fig. 5C**). Our data were further strengthened by rescue experiments where we co-expressed *UAS-CNMa* in *3.1Lsp2>Perk*^*RNAi*^ or *>Gcn2*^*RNAi*^ animals. Strikingly, we observed a near-complete rescue of the egg retention phenotypes (**Fig. 5F-K)**. Based on these data, we conclude that Atf4-mediated transcriptional regulation of *CNMa* in the fat body regulates ovulation.

Finally, we sought to identify which neurons respond to CNMa to promote egg-laying behavior. In *Drosophila*, egg-laying behavior is controlled by several groups of neurons, including two pairs of neurons descending from the mushroom body called oviposition descending neurons (oviDNs)^37^. The oviDNs are a subset of sexually dimorphic neurons in which the *P1* promoter of *fruitless (fru*^*P1*^) is active^37^. Synaptic silencing of all *fru*^*P1*^-expressing cells blocks ovulation, as does ablation of oviDNs^37^. We tested whether the CNMa receptor, CNMaR, is broadly required in all *fru*^*P1*^-expressing neurons; however, RNAi depletion of *CNMaR* using *fru*^*P1*^*-GAL4* resulted in an apparent block in oogenesis (**Fig. S4C-E**), which likely would obscure an egg-retention phenotype. To circumvent this, we next depleted CNMaR specifically in the oviDNs using a *split-GAL4* driver combination^37^ (abbreviated here as *oviDN-GAL4*). Consistent with our hypothesis that CNMa from the fat body signals to neurons that promote ovulation, we saw that RNAi-mediated depletion of *CNMaR* using an *oviDN*-*GAL4* resulted in an increased number of eggs retained per ovary compared with control animals (**Fig. 5L**; 29.19 ± 1.45 in *oviDN>CNMaR*^*RNAi*^ vs. 24.17 ± 2.16 in control; p<0.05).

Taken together, these findings support a model whereby ISR signaling in the fat body induces CNMa production, which is secreted to activate ovulation-promoting neuronal circuits, including at least the oviDNs.

## Discussion

Single-celled organisms like yeast, where Atf4 (Gcn4 in yeast) was first discovered^43,44^, rely on the ISR pathway under conditions of nutrient deprivation rather than for survival. In contrast, metabolically active tissues in higher organisms have evolved to rely on the ISR pathway for their homeostatic function^45^. This is illustrated both by constitutive activity of Atf4 in fat tissues and by the metabolic phenotypes seen in *Atf4* mutant *Drosophila* and mice^4,7^. An increasing body of literature supports a role for fat tissues in modulating peripheral organ function^46^, but whether and how this is regulated by ISR signaling is under active investigation. Our study makes a significant dent in this open problem by demonstrating multiple mechanisms by which homeostatic ISR signaling informs oogenesis in female flies.

Our findings show a requirement for both known ISR kinases, Perk and Gcn2, in both oogenesis and ovulation. Given that Perk is sensitive to changes in lipid content^47^ and Gcn2 senses amino acid content^48^, we favor a model whereby ISR signaling in the fat body acts as a conduit between nutrient status and reproductive capacity (**Fig. 6**). In one branch of this model, oocyte maturation is compromised in the absence of Atf4 due to reduced yolk lipoprotein trafficking from the fat body to the ovary. In another seemingly parallel branch of this model, ISR signaling regulates ovulation, which is further downstream of oogenesis, by neuromodulation of sexually dimorphic neurons. The sum effect of these two regulatory branches likely leads to the observed follicle death and oogenesis arrest observed in ISR-deficient animals, and manifests as an overall decrease in fertility seen in global *Atf4* mutants^18^. These findings suggest that Atf4 signaling in the fat body regulates parallel processes with synergistic effects on female reproductive output.

**Figure 6.**
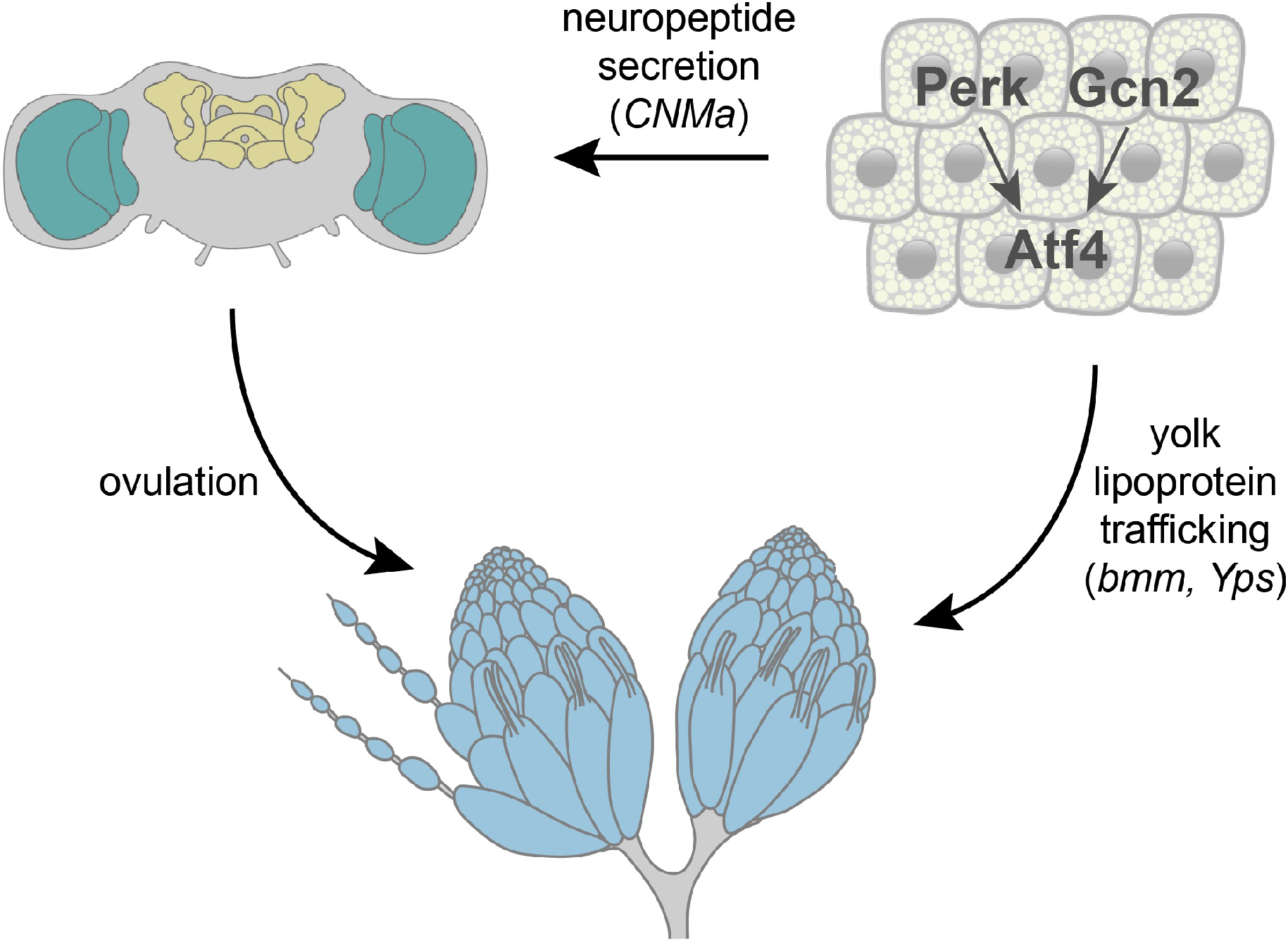
Model describing a role for ISR signaling in inter-organ regulation of reproduction.

### ISR signaling in the fat body supports oogenesis non-autonomously via yolk trafficking

The loss of oocyte yolk granule abundance we saw in the absence of *Atf4* (**Fig. 2E-F**) could be due to impaired lipoprotein synthesis, assembly, or secretion, causing less yolk to successfully reach maturing oocytes. Our results suggest that the decreased yolk abundance in mature oocytes is likely due to a combination of reduced yolk protein synthesis and impaired lipid droplet formation (**Fig. 2G-J**). The dramatic reduction in Yp1::GFP abundance in fat seen with loss of *Atf4* (**Fig. 2G-H**) can be attributed to reduced yolk protein transcript (**Fig. 2K**), though it remains unclear whether this is because of direct transcriptional regulation of *Yp1-3* by Atf4 or secondary effects of loss of *Atf4* in the fat body.

Lipoprotein assembly relies on mobilization of triacylglycerol (TAG) reserves from lipid droplets, which has been demonstrated to be regulated by Bmm^32^. Our data found that fat bodies lacking *Atf4* have reduced levels of *bmm* transcripts (**Fig. 2K**), which likely yields the defects in neutral lipid distribution we observed in these adipocytes (**Fig. 2I-J**). These findings are consistent with a previous report that TAG breakdown is compromised in *Atf4* mutants^4^. Additionally, since lipid droplets form from extensions of the ER membrane, it is also possible that structural changes to the ER may occur upon loss of *Atf4* in fat tissue. This possibility is supported by our preliminary data where we see marked differences in fat body ER structures between control and *3.1Lsp2>Atf4*^*RNAi*^ animals (**Fig. S5)**. However, further investigation is needed to determine if these changes to ER structures are *bmm*-dependent. In either case, loss of lipid homeostasis in the fat body can impact vitellogenesis in at least two different ways: first, the loss of lipid droplets decreases the yolk lipoprotein trafficking to the ovary; second, lipid droplets serve to shuttle cholesterol-derived molecules such as the steroid hormone ecdysone. Ecdysone signaling has been shown to promote vitellogenic events such as the accumulation of lipids in the maturing oocyte^49^. The loss of lipid droplets in the absence of *Atf4* may attenuate such pro-vitellogenic signals, which would result in the observed reduction in oocyte yolk granules. This supposition also lends an explanation for how the fat body senses yolk lipoprotein needs of the ovary. The ovary is known to be a major source of ecdysone in adult females, with ecdysone synthesis beginning in stage 9 follicles^50^. This leads us to propose ecdysone as a reasonable candidate for a feedback signal that might communicate ovary yolk lipoprotein needs to the fat body. Our future work will address if the ovary-derived steroid hormone interacts with ISR signaling in the fat body to stimulate yolk production.

### The role of ISR signaling in regulating oogenesis under stress

Our results demonstrate that Atf4 activity in the fat body is required for oogenesis recovery after acute amino acid deprivation (**Fig. 3C, F, G**), positioning it as a metabolic sensor that restricts or permits oogenesis based on nutrient availability. The increased follicle death we observed in *3.1Lsp2>Atf4*^*RNAi*^ ovaries from “re-fed” females (**Fig. 3A**) overwhelmingly occurred during mid-oogenesis (**Fig. 3G**), which coincides with the stage when yolk lipoprotein uptake by the oocyte commences. We found that yolk lipoprotein trafficking to the ovary is Atf4-dependent (**Fig. 2E-L**). Thus, it is possible that yolk lipoprotein trafficking from the fat body to the ovary is involved in relaying nutrient status to the ovary downstream of Atf4. This possibility is supported by our preliminary data showing that depleting *Yp1* in the fat body is sufficient to cause increased follicle death (**Fig. S2D**).

Atf4 is basally and constitutively active in the adult fat body^4,7^, making it poised to support reproduction under homeostatic conditions. However, Atf4 translation also increases in response to stress, notably amino acid deprivation, in the fat body and other tissues^7,51^. Amino acid deprivation has a predictably negative impact on fertility, accompanied with a reduction in germline stem cells of the ovary, which has been previously demonstrated to be dependent on fat body Gcn2 signaling^52^. How Atf4 signaling in the fat body differs between homeostatic vs. amino acid deprivation conditions remains an open question. Atf4 is a member of the ATF/CREB family of bZIP transcription factors with known interacting partners^53,54^. Thus, an attractive possibility is that Atf4 regulates target gene expression in the fat body in concert with co-factors which change under different environmental conditions. Ongoing work aims to dissect the molecular mechanisms by which Atf4 and ISR signaling regulate fertility changes under homeostatic and stress conditions.

### ISR signaling in the fat body promotes ovulation via CNMa secretion

Loss of either ISR kinase resulted in increased follicle death, mid-oogenesis arrest, and reduced yolk granules in maturing oocytes (**Fig. 4A-E)**, demonstrating that both Perk and Gcn2 can act upstream of Atf4 in the adult fat body. Further, we observed that fat body knockdown of *Perk* or *Gcn2* resulted in retention of mature oocytes in the ovary (**Fig. 4H, I**), which is indicative of a defect in ovulation. We reason that depletion of either ISR kinase results in milder loss of Atf4, resulting in only a partial loss of ISR signaling, as opposed to *Atf4* knockdown which results in greater loss of ISR signaling. Substantial loss of ISR signaling leads to a concomitantly substantial oogenesis defect, thus obscuring any ovulation defects (**Fig. 4G, K**). This notion is further supported by our data showing that joint knockdown of *Perk* and *Gcn2*, which leads to greater loss of ISR signaling in the fat body, does not display ovulation defects (**Fig. S3D**).

Our data demonstrate that ISR directly mediates ovulation via multiple neuropeptides, including CNMa and Eth (**Fig. 5A,B**), with CNMa being the largest contributor to ovulation. Based on our analysis, we propose that CNMa is likely transcriptionally regulated by Atf4 in the fat body (**Fig. 5C, S2A**). This is supported by our finding that *CNMa* is constitutively expressed in larval and adult fat body under homeostatic conditions, similar to Atf4 (**Fig. S5C)**. Intriguingly, recent work has implicated CNMa in the amino acid deprivation response downstream of Gcn2 in enterocytes^42^. However, this study utilized pan-neuronal drivers to demonstrate the role of CNMa-CNMaR signaling axis in feeding behavior. In the context of ovulation, our data show that CNMa signaling via CNMaR appears to activate neurons such as the oviDNs that are known to promote egg-laying behavior^37^. It is worth noting that CNMaR silencing specifically in oviDNs caused egg retention to a lesser degree than loss of CNMa from the fat body (**Fig. 5L**). This suggests that other cells (in addition to the oviDNs) may be responsive to CNMa in the context of egg-laying, including neurons in the central or peripheral nervous systems. For example, octopaminergic neurons in the brain and reproductive tract promote ovulation steps in response to mating-induced octopamine^38^. Additionally, peripheral neurons that innervate the reproductive tract have been shown to regulate ovulation^40^. High-throughput expression data generated by the modENCODE^30,31^ and FlyAtlas^55,56^ projects suggest that *CNMaR* expression is restricted to the head region, though it is possible that there are low levels of *CNMaR* in other cells. Further studies are needed to identify the entire cohort of cell types that are CNMa-responsive in the regulation of ovulation downstream of ISR signaling in the fat.

## Methods

### Fly husbandry and stocks

The transgenic lines used in this study are publicly available via Bloomington *Drosophila* Stock Center or were sourced from other labs. See **Table S1** for a complete list of lines used.

Animals were cultured in standard cornmeal agar media containing yeast and molasses (LabExpress, Inc). Fly stocks were maintained at either room temperature (RT) or 18°C and experimental crosses were maintained at 25°C. Conditional expression studies utilizing the *GAL80*^*TS*^ transgene were maintained at 18°C to restrict GAL4 activity, and adult progeny were shifted to 25°C to permit GAL4-mediated knockdown and/or over-expression. Adult females were collected at 0-2 days following eclosion, mated with *yw* males in standard media supplemented with yeast, and dissected at the indicated ages. For starvation assays, mated females were raised on amino acid-deficient medium containing 5% sucrose in 2% agar for four days. To re-feed, females were transferred to fresh yeast-molasses medium vials for two more days.

### Immunofluorescence

Adult ovaries and larval/adult fat body were dissected in PBS and fixed in 4% PFA for 15 minutes at RT. Samples were washed in PBS+0.1% detergent (Triton X-100 for ovary, Tween-20 for fat body) twice and incubated overnight at 4°C in primary antibody solution. Secondary antibodies along with DAPI (300nM final concentration) were incubated for 2 hours at RT in the dark. Guinea pig anti-Tj (M. Van Doren) was used at 1:10,000. Mouse anti-KDEL (Santa Cruz Biotech) was used at 1:100. Phalloidin-Rhodamine (Life Technologies) was used at 1:500. For yolk granule visualization, ovaries were fixed, washed, and mounted without DAPI. Yolk granules were visualized by auto-fluorescence upon excitation by 405nm laser^57^. For neutral lipid staining in the adult fat body, samples were fixed and washed, followed by incubation in BODIPY solution (Thermo Scientific) for 20 minutes at a final concentration of 2 µg/mL. Confocal images were captured using a Nikon A1 confocal microscope through the Center for Biological Imaging at the University of Pittsburgh. Egg retention representative images were captured using a Nikon SMZ1270I with a ring light and DS-Ri2 camera attachment.

### Fluorescent in situ hybridization (FISH)

FISH detection of *Atf4* mRNA was performed using a commercially available kit (Molecular Instruments, Inc) and was conducted as per manufacturer’s instructions.

### ChIP-seq data analysis via ENCODE

ChIP-seq was performed on Atf4-GFP by Dr. Kevin White (UChicago) and is publicly accessible via the ENCODE^30,58^ project website (www.encodeproject.org). ENCODE accession number for dataset used: ENCFF986LGA. Briefly, embryos (0-22h) were isolated from *crc-MiMIC-GFP* transgenic fly line and ChIP was performed using an anti-GFP antibody with sequencing on the Illumina HiSeq 2000 platform. Reads were aligned to the dm6 *Drosophila melanogaster* genome annotation. Read frequency plots shown in **Fig. S2A** were generated by loading ENCODE bigWig file onto the Integrated Genome Viewer software (www.igv.org; Broad Institute).

### Quantitative RT-PCR

RNA was isolated from four female adults per replicate using TRIzol (Invitrogen) according to manufacturer’s protocol. Reverse transcription was performed using Maxima H minus reverse transcriptase (Thermo Scientific) according to manufacturer’s protocol. qPCR amplification reactions were prepared using SYBR Green Master Mix (Thermo Scientific) in the BioRad CFX96 real time system. See **Table S3** for list of primers used for analysis.

### Quantifications and statistical analyses

To quantify border cell migration, area and distance measurements were determined by ImageJ analysis. For egg retention quantifications, ovaries were dissected with minimal perturbance, fixed, washed, and mounted. The number of mature oocytes per ovary was then scored manually. The appropriate statistical analyses for each assay (indicated in the accompanying legend) were performed using Microsoft Excel or GraphPad Prism.

## Figure legends and tables

**Figure S1.**
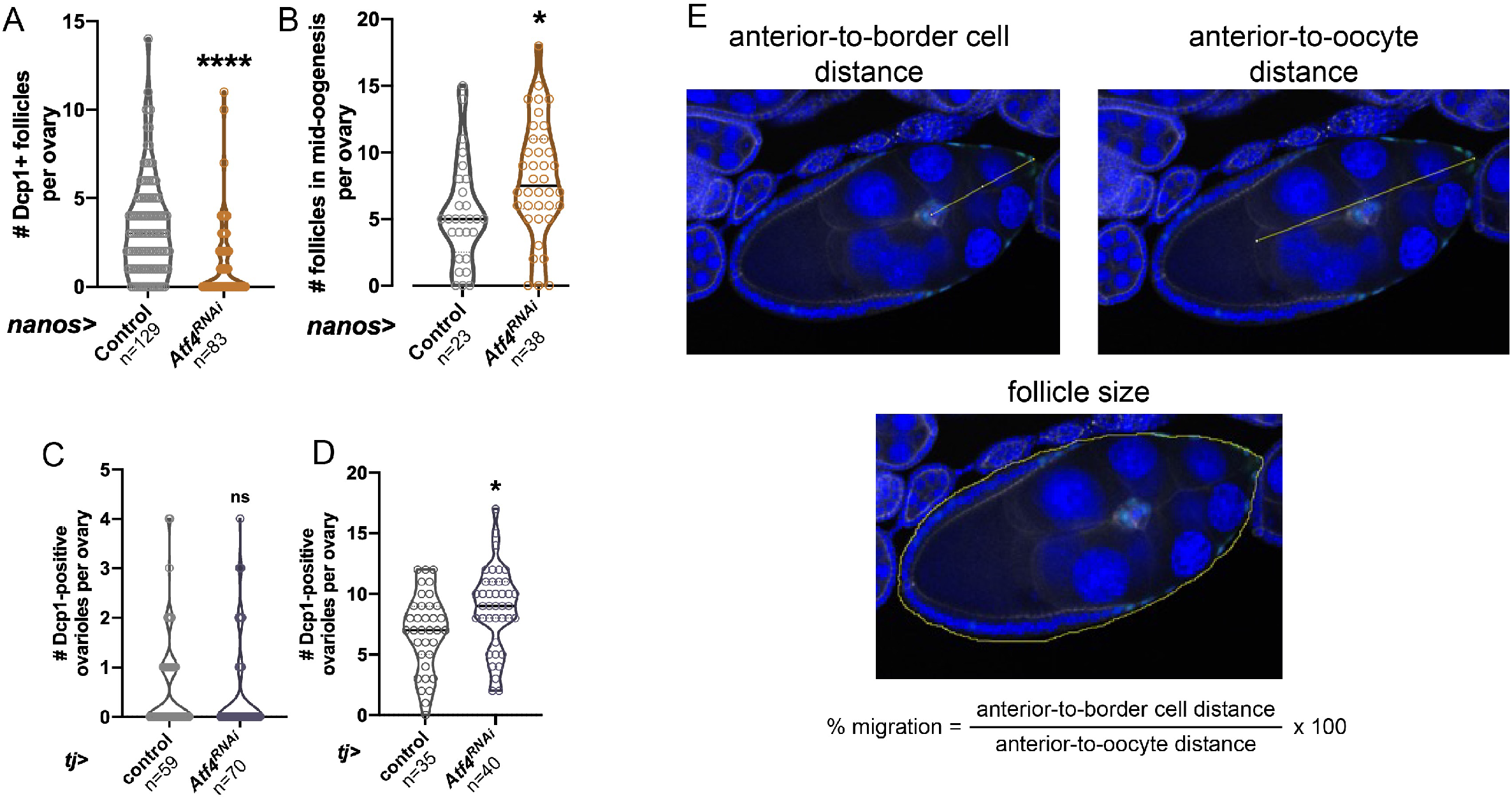
(A-B) Quantification of follicle death as determined by Dcp1 staining (A) or the number of follicles in mid-oogenesis (B) from 3-day old adult females in control (*lacZ*) or *Atf4* depletion using the germline driver *nanos-GAL4*. (C-D) Quantification of follicle death as determined by Dcp1 staining (C) or the number of follicles in mid-oogenesis (D) from 3-day old adult females in control (*lacZ*) or *Atf4* depletion using the somatic driver *tj-GAL4*. (E) Illustrative details of quantification of border cell migration shown in **Fig. 1F**. Distance and area values were calculated using ImageJ.

**Figure S2.**
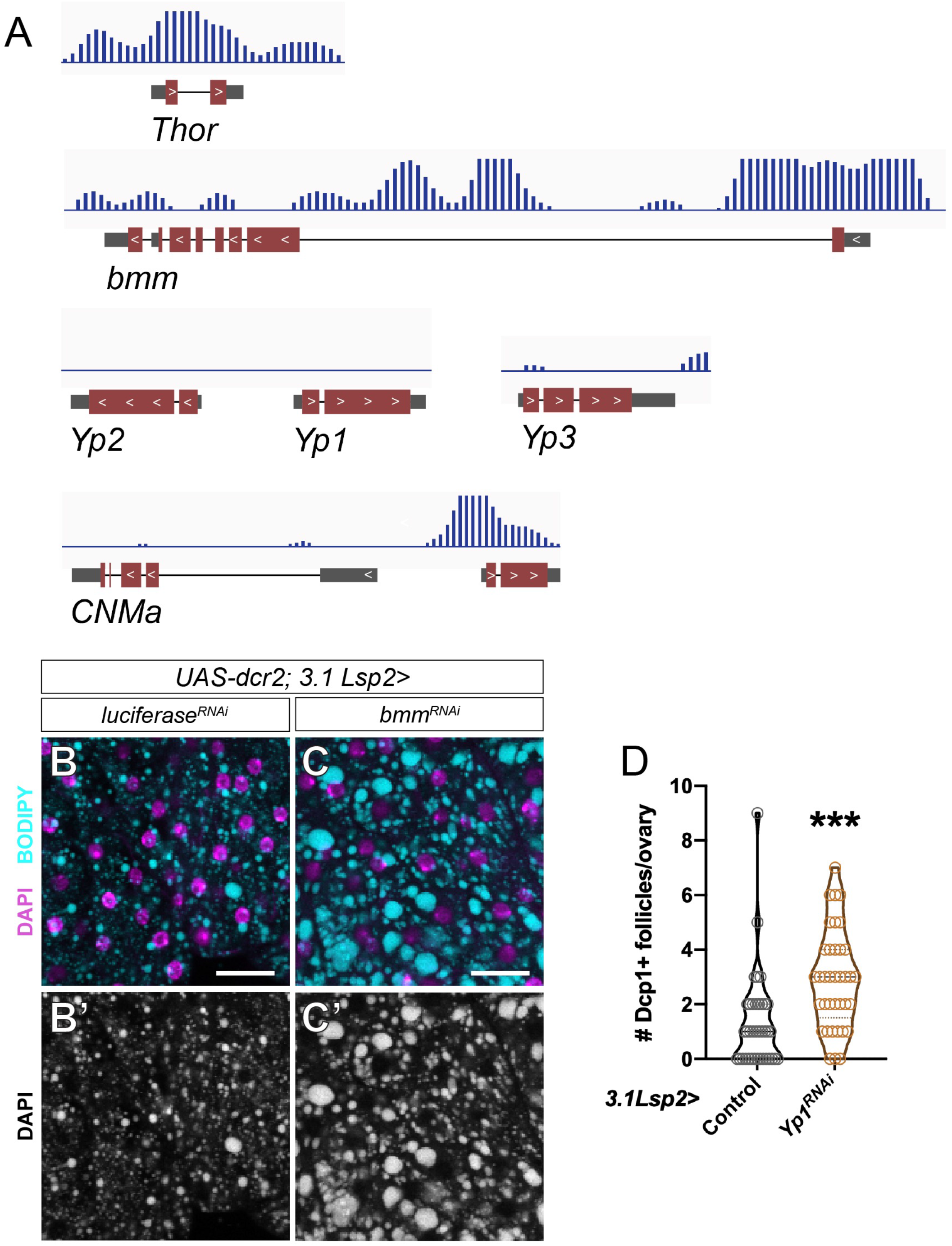
(A) Atf4 occupancy read abundance at the indicated loci based on previously published ChIP-seq analysis of Atf4-GFP in *Drosophila* embryos^30,31^. In gene schematics, gray boxes represent UTRs, red boxes represent coding exons, and black lines represent introns. (B-C) Representative confocal images of neutral lipid staining (BODIPY, cyan) in adult fat body following control (*luciferase*, B) or *bmm* (C) depletion using the fat body-specific driver *3.1Lsp2-GAL4. UAS-dcr2* was added to both genotypes to improve the RNAi efficiency. Scale bar: 25 µm. (D) Quantification of follicle death following depletion of *Yp1* from the fat body using *3.1Lsp2-GAL4*.

**Figure S3.**
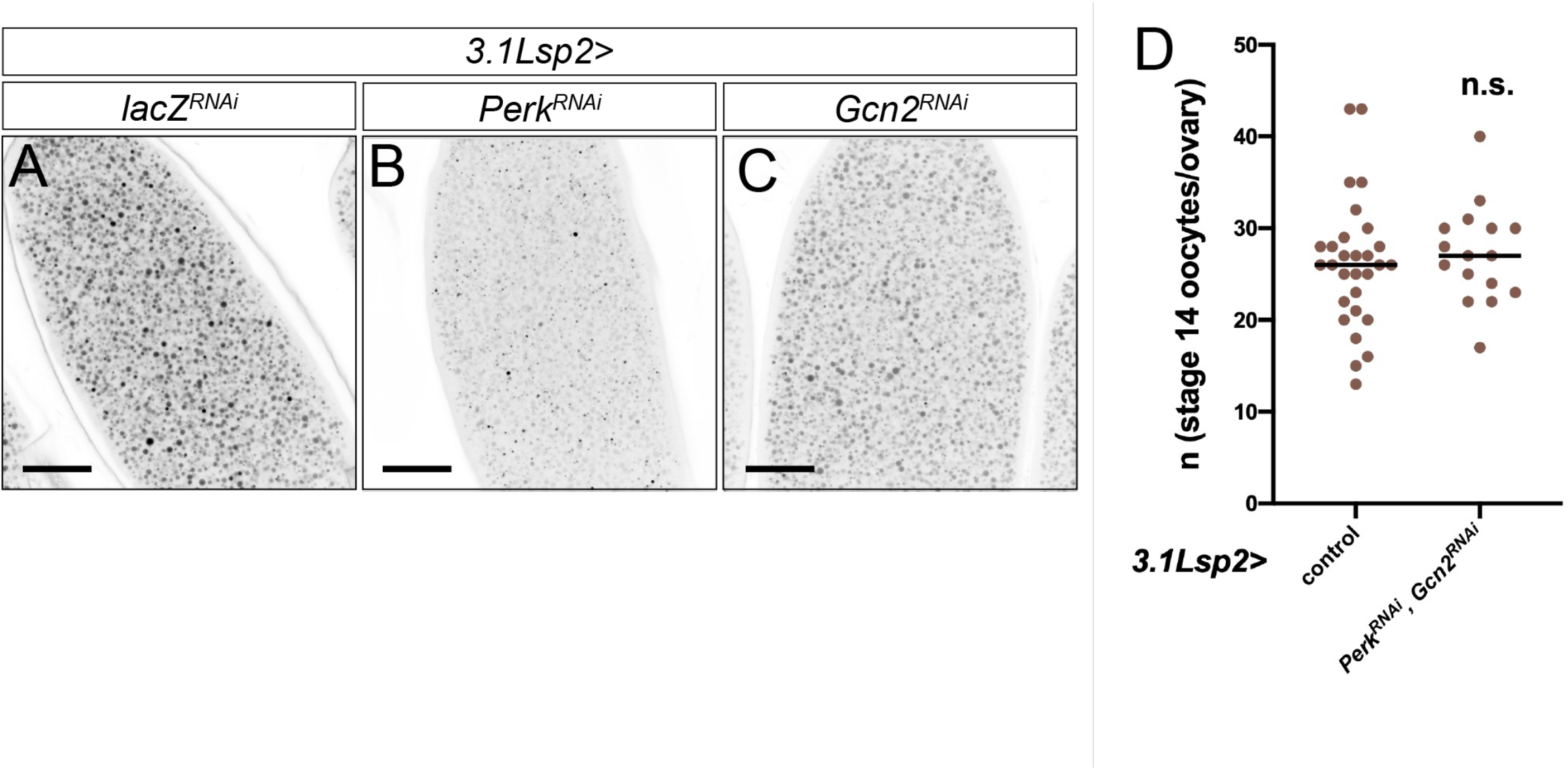
(A-C) Representative confocal images of stage 14 oocytes from *3.1Lsp2>lacZ*^*RNAi*^ (A), *Perk*^*RNAi*^ (B), and *Gcn2*^*RNAi*^ (C) females using 405nm laser to visualize yolk granules. Scale bar: 200 µm. (D) Quantification of oocytes per ovary upon double knockdown of *Perk* and *Gcn2* in fat tissue using *3.1Lsp2-GAL4* compared with *control*^*RNAi*^.

**Figure S4.**
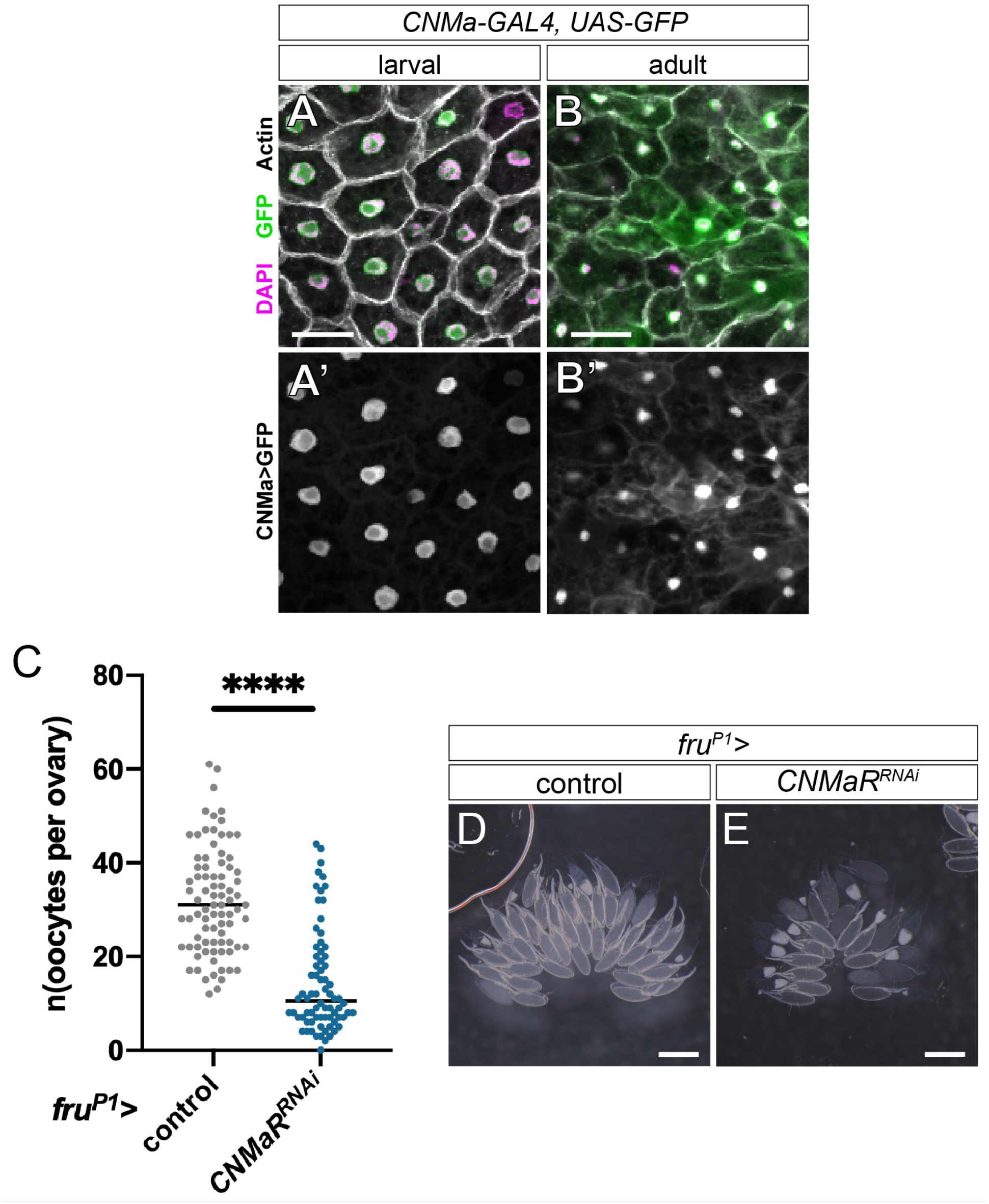
(A-B) Visualization of a CNMa-GAL4 driver in larval (C) and adult (D) fat tissues using UAS-GFP. (C) Quantification of oocytes contained per ovary following control or *CNMaR* depletion in *fru*^*P1*^-expressing cells. Black lines denote average for each genotype. (D-E) Representative ovaries following control (D) or *CNMaR* depletion (E) in *fru*^*P1*^-expressing cells. Scale bar: 500 µm.

**Figure S5.**
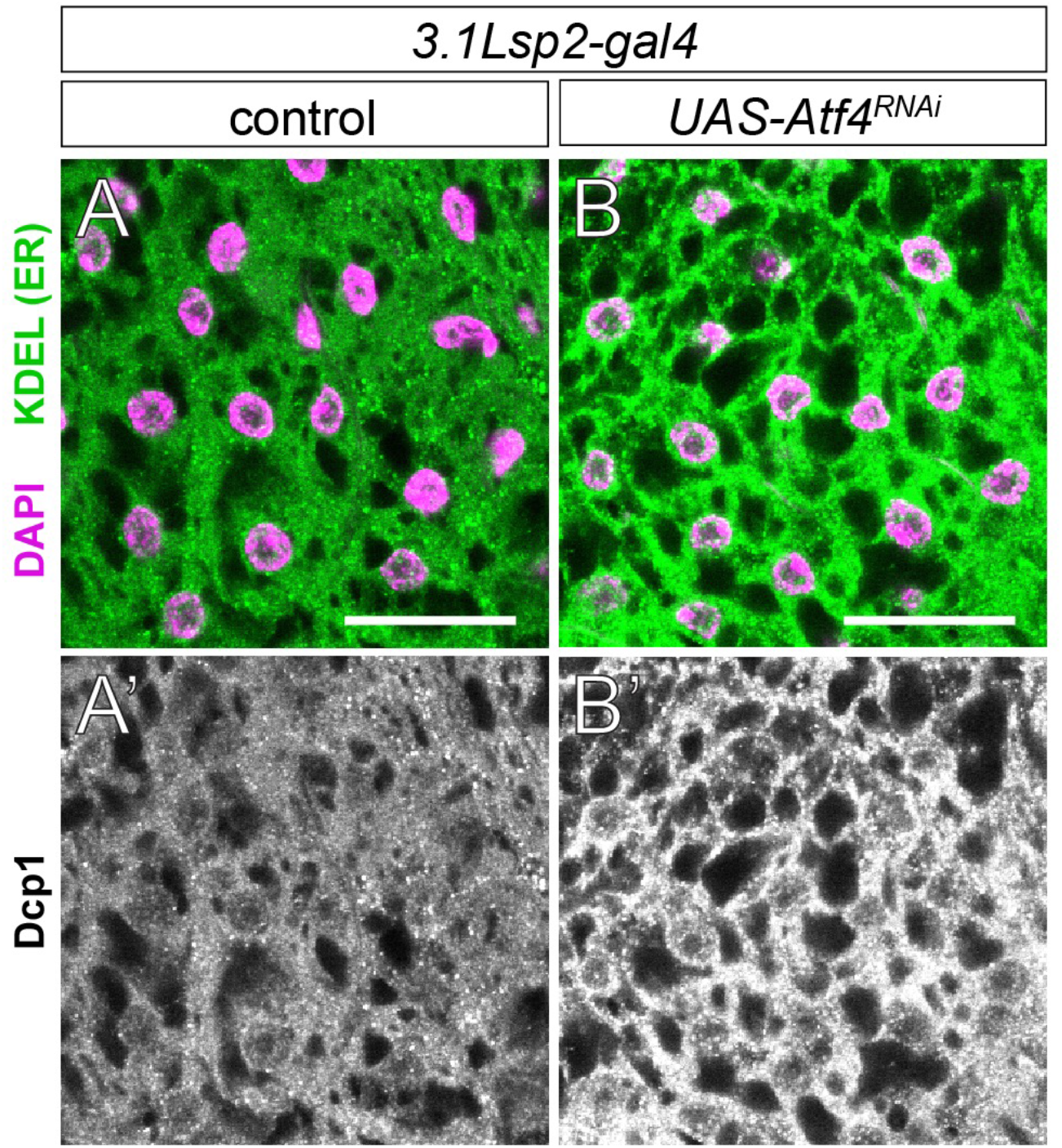
(A-B) ER labeling of representative control (a) and *3.1Lsp2>Atf4*^*RNAi*^ (b) fat tissues using anti-KDEL antibody (green). DAPI (magenta) labels nuclei. Scale bar: 25 µm.

**Table S1.** List of transgenic and mutant fly lines used in this study with stock numbers or source as indicated.

| Genotype | Source | Figures used |
| --- | --- | --- |
| *P{GawB}tj<sup>NP1624</sup> (tj-GAL4)* | Kyoto #104055 | 1D-F; S1E |
| *nanos-gal4::VP16* | BDSC #4937 | S1A-B |
| *3.1Lsp2-GAL4* | BDSC #84285 | 2A-J,L; 3B-G; 4A-K; 5A-B,D-K; S3 |
| *oviDN-GAL4* | BDSC #86832 | 5L |
| *fru<sup>P1</sup>-GAL4* | BDSC #66696 | S4C-E |
| *tub-GAL80<sup>TS</sup>* | BDSC #7019 | 4F-K; 5D-K |
| *yw* | BDSC #1495 | 1B-C; 2K; 5C |
| *crc*<sup>GFSTF</sup> (MI02300-GFSTF) | BDSC #59608 | 2K; 5C |
| *Yp1::GFP* | Dr. Yusuke Hara | 2G-H |
| *UAS-lacZ*<sup>RNAi</sup> | not publicly available | 2A, E, G, I; 4A; 5B; S3A |
| *VIE-260B* (control; isogenic background control for *Atf4/Perk/Gcn2*<sup>RNAi</sup>) | VDRC #106174 | 1D, F; 2C, D, L; 3B-E; 4D-F, K; 5D, F, I; S1A-D; S2D; S3D; S4C, D; S5 |
| *UAS-luciferase*<sup>RNAi</sup> | BDSC #31603 | 5A; S2B |
| *UAS-Atf4*<sup>RNAi</sup> | VDRC #109014 | 1E,F; 2B-D,F,H,J,L; 3B-C,F-G; 4D,G, K; S1A-D; S5B |
| *UAS-Atf4*<sup>RNAi</sup> (2) | VDRC #2934 | 1F |
| *UAS-Perk*<sup>RNAi</sup> | VDRC #110278 | 4B,D-E,H, K; 5G, J; S3B, D |
| *UAS-Gcn2*<sup>RNAi</sup> | VDRC #103976 | 4C-E, I, K; 5H, K; S3C-D |
| *UAS-CNMa*<sup>RNAi</sup> | VDRC #110796 | 5B, E |
| *UAS-ETH*<sup>dsRNA</sup> | BDSC #26242 | 5A |
| *UAS-sNPF*<sup>RNAi</sup> | BDSC #25867 | 5A |
| *UAS-Tk*<sup>RNAi</sup> | BDSC #25800 | 5A |
| *UAS-CNMaR*<sup>RNAi</sup> | VDRC #101076 | 5L; S4C, E |
| *UAS-Yp1*<sup>RNAi</sup> | BDSC #67219 | S2D |
| *UAS-bmm*<sup>RNAi</sup> | BDSC #25926 | S2C |
| *UAS-dcr2* | BDSC #24651 | 5A; S2B-C |
| *UAS-bmm* | BDSC #76600 | 2L |
| *UAS-TetxLC.tnt* (*UAS-TnT*) | BDSC #28837 | 4J-K |
| *UAS-CNMa* | Dr. Won-Jae Lee | 5I-K |

**Table S2.** List of *Drosophila* neuropeptides and their gene symbols. Candidates were included if they bore the “neuropeptide” classification on Flybase, and this list was compared to a previously curated list of neuropeptides^59^ to verify that none were missing. Genes highlighted in green indicate loci in which Atf4 occupancy was determined by ChIP-seq^30,31^.

| Neuropeptide gene | Symbol | Neuropeptide gene | Symbol |
| --- | --- | --- | --- |
| *Adipokinetic hormone* | *Akh* | *Insulin-like peptide 5* | *dilp5* |
| *amnesiac* | *amn* | *Insulin-like peptide 6* | *dilp6* |
| *Allatostatin A* | *AstA* | *Insulin-like peptide 7* | *dilp7* |
| *Allatostatin C* | *AstC* | *Insulin-like peptide 8* | *dilp8* |
| *Allatostatin double C* | *AstCC* | *ion transport peptide* | *ITP* |
| *Bursicon* | *Burs* | *Leucokinin* | *Lk* |
| *Capability* | *Capa* | *Myoinhibiting peptide precursor* | *Mip* |
| *Crustacean cardioactive peptide* | *CCAP* | *Myosuppressin* | *Ms* |
| *CCHamide-1* | *CCHa1* | *neuropeptide F* | *NPF* |
| *CCHamide-2* | *CCHa2* | *Neuropeptide-like precursor 1* | *Nplp1* |
| *CNMamide* | *CNMa* | *Neuropeptide-like precursor 2* | *Nplp2* |
| *Corazonin* | *Crz* | *Neuropeptide-like precursor 3* | *Nplp3* |
| *Diuretic hormone 31* | *Dh31* | *Neuropeptide-like precursor 4* | *Nplp4* |
| *Diuretic hormone 44* | *Dh44* | *Orcokinin* | *Orcokinin* |
| *Drosulfakinin* | *Dsk* | *Partner of Bursicon* | *Pburs* |
| *Ecdysis triggering hormone* | *Eth* | *Pigment-dispersing factor* | *Pdf* |
| *Eclosion hormone* | *Eh* | *Proctolin* | *Proc* |
| *FMRFamide* | *FMRFa* | *Prothoracicotrophic hormone* | *PTTH* |
| *Glycoprotein hormone alpha 2* | *Gpa2* | *RYamide* | *RYa* |
| *Glycoprotein hormone beta 5* | *Gpb5* | *SIFamide* | *SIFa* |
| *Hugin* | *Hug* | *short neuropeptide F precursor* | *sNPF* |
| *Insulin-like peptide 1* | *dilp1* | *Sex Peptide* | *SP* |
| *Insulin-like peptide 2* | *dilp2* | *space blanket* | *spab* |
| *Insulin-like peptide 3* | *dilp3* | *Tachykinin* | *Tk* |
| *Insulin-like peptide 4* | *dilp4* |  |  |

**Table S3.**
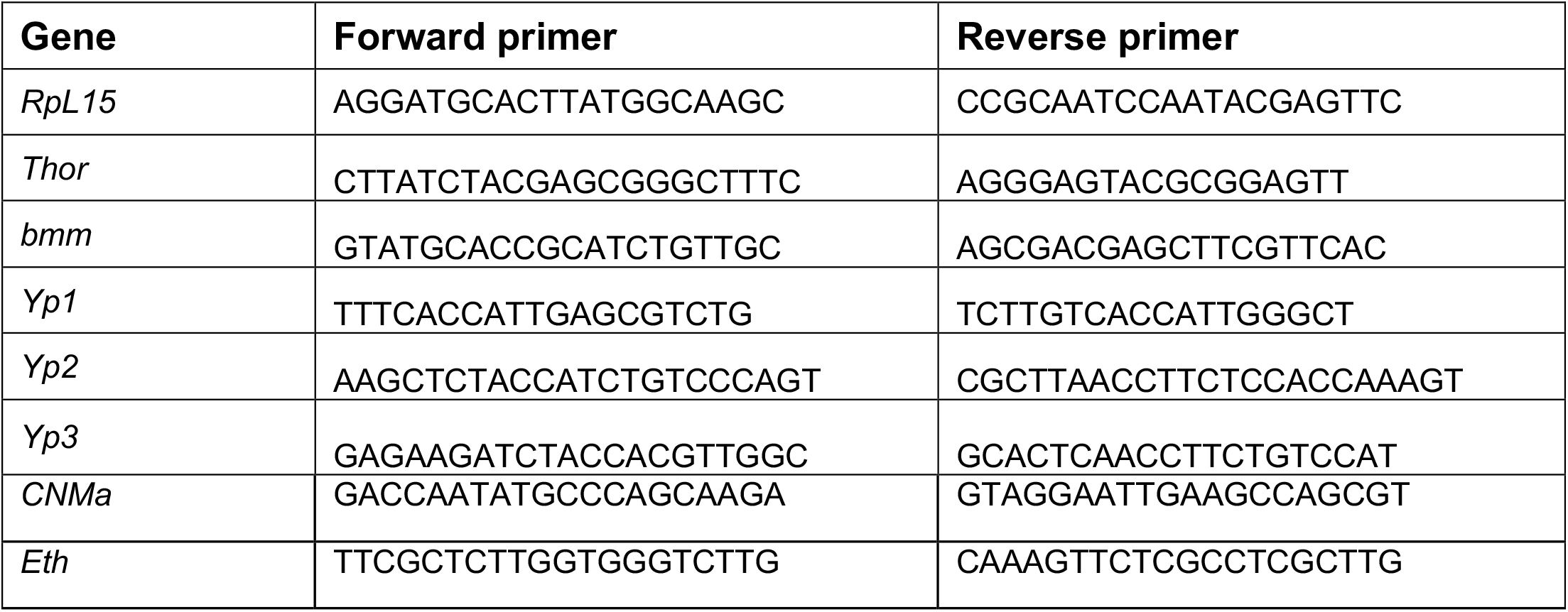
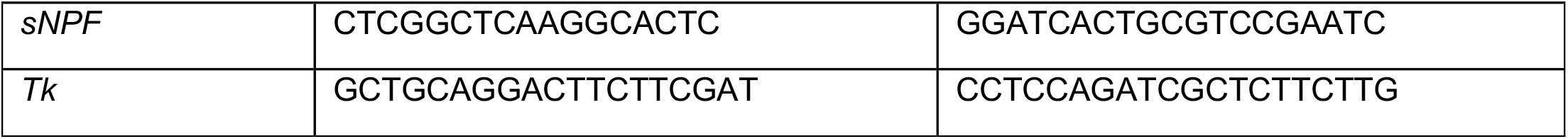
Primers used for qPCR analysis in this study are listed below.

## Acknowledgements

We are deeply grateful to the Center for Biological Imaging at the University of Pittsburgh for imaging assistance and access to equipment. Additionally, we thank the Bloomington *Drosophila* Stock Center (BDSC, Bloomington, IN), the Developmental Studies Hybridoma Bank (DSHB, Iowa City, IA), the Vienna *Drosophila* Resource Center (VDRC, Vienna, Austria), and the Transgenic RNAi Project (Harvard Medical School, Cambridge, MA) for making available reagents needed for this study. We are grateful to Drs. Marc Amoyel, Elizabeth Ables and Shyama Nandakumar for providing project and manuscript feedback. We also thank Drs. Won-Jae Lee and Yusuke Hara for generously sharing transgenic fly lines. D.V. is funded by NIH R00EY029013 and L.G. is funded by NIH T32DK063922 and NIH K99GM149982.

## Author contributions

Conceptualization: L.G. and D.V.; Data curation: L.G., M.M., E.L., N.N.V.P., and D.V.; Formal analysis: L.G., M.M, and D.V.; Validation: L.G., M.M., E.L., N.N.V.P., and D.V.; Methodology: L.G and D.V.; Writing – original draft: L.G. and D.V.; Writing – review and editing: L.G., M.M., and D.V.; Supervision: L.G. and D.V.

